# Cell Chirality of Micropatterned Endometrial Microvascular Endothelial Cells

**DOI:** 10.1101/2023.10.20.563368

**Authors:** Samantha G. Zambuto, Ishita Jain, Hannah S. Theriault, Gregory H. Underhill, Brendan A.C. Harley

## Abstract

Chirality is an intrinsic cellular property that describes cell polarization biases along the left-right axis, apicobasal axis, or front-rear axes. Cell chirality plays a significant role in the arrangement of organs in the body as well as the orientation of organelles, cytoskeletons, and cells. Vascular networks within the endometrium, the mucosal inner lining of the uterus, commonly display spiral architectures that rapidly form across the menstrual cycle. Herein, we systematically examine the role of endometrial-relevant extracellular matrix stiffness, composition, and soluble signals on endometrial endothelial cell chirality using a high-throughput microarray. Endometrial endothelial cells display marked patterns of chirality as individual cells and as cohorts in response to substrate stiffness and environmental cues. Vascular networks formed from endometrial endothelial cells also display shifts in chirality as a function of exogenous hormones. Changes in cellular-scale chirality correlate with changes in vascular network parameters, suggesting a critical role for cellular chirality in directing endometrial vessel network organization.

## 1. Introduction

The endometrium is the mucosal lining of the uterus that consists of a functionalis and a basalis layer^1^. The functionalis responds to hormones and proliferates, differentiates, and sheds based on the hormonal signals it receives whereas the basalis does not^1^. During the menstrual cycle, the endometrium undergoes cyclic vascularization and angiogenesis orchestrated by fluctuating levels of steroidal sex hormones^2^, notably 17β-estradiol (E2) and progesterone (P4)^3^. Here, the proliferative phase is often characterized by E2-dominated proliferative growth^3^. The subsequent secretory phase is largely a P4-dominated phase that instructs endometrial priming and differentiation^3^. In the absence of an implanted blastocyst, the menstrual phase is marked by P4 withdrawal and subsequent tissue edema, vessel permeability, an influx of leukocytes, and increased blood flow associated with endometrial shedding and menses^3^.

The endometrium is a rare adult human tissue capable of non-pathological, rapid angiogenesis, making it a model system for understanding functional vascular remodeling. The endometrial vasculature also plays a key role in a variety of disorders, including preeclampsia, recurrent miscarriage, and failed implantation during pregnancy^2, 4^. Here, additional hormones and biomolecules may act in concert with or against steroidal sex hormones to perturb the endometrial vasculature. For example, glucocorticoids such as cortisol, steroid hormones produced in response to stress by the adrenal cortex,^5^ can have a detrimental effect on embryo implantation, pregnancy outcomes, and fetal health ^6–8, 9^. Elevated cortisol has previously been shown to reduce endothelial cell tube formation and trophoblast invasion in two-dimensional assays^7, 10^. Defining how cortisol affects endometrial cells may lead to improvements in maternal and fetal outcomes and offers exciting potential to incorporate a new class of biomolecular signal into tissue engineering models of the endometrium and trophoblast invasion.

Chirality is an essential design concept across biological systems. Chirality influences arrangement of organs and tissues, the right-handed bias in twining plants, and the adaptation of chiral morphology in bacterial colonies under stress^11, 12^. Chirality at the cellular level has been observed as directional rotation associated with organelles, the cytoskeleton, and even cells themselves^12^. Differences in cell chirality persist as a function of phenotype and between different cell types such as recent observation of an anti-clockwise bias in skeletal muscle cells but a clockwise bias in endothelial cells in micropatterning systems^12^. In the context of cell cohorts, changes in cell chirality have been shown to disrupt endothelial cell permeability^13^. And while the endometrium is marked by a characteristic spiral organization of endometrial vascular structures^2^, the role of the endometrial tissue microenvironment on endothelial cell chirality remains unclear^13^.

Herein, we systematically examine the combined role of endometrial-associated extracellular matrix stiffness, extracellular matrix ligand presentation, and angiogenic (VEGF) vs. stress-associated (cortisol) biomolecules on endometrial endothelial cell chirality. First, we define patterns of endometrial endothelial cell attachment and chirality in response to single and pairwise combination of 10 endometrial-associated^14^ extracellular matrix (ECM) ligands using a high-throughput 2D array format. We report shifts in endometrial endothelial cells chirality using three separate metrics to describe polarization biases along one of three axes:^13^ the left-right axis, apicobasal axis, or front-rear axis of the cell. We report 3 distinct chirality phenotypes of endometrial endothelial cells as a function of their underlying ECM environment via principal component analyses. Representative ECM combinations from these 3 chirality clusters were subsequently used to evaluate cell chirality sensitivity to exogenous VEGF and as a function of cortisol dosage. Finally, we link changes in cell chirality on 2D extracellular matrix functionalized substrates to shifts in the structure and organization of three-dimensional vascular networks formed using endometrial endothelial cells in response to a panel of stress and decidualization hormones. Shifts in endometrial vascular network parameters are correlated with changes in cellular level chirality metrics, suggested a key role for cellular chirality in directing the etiology of vessel network architectures.

## 2. Results

### 2.1. Human endometrial microvascular endothelial cells display differential attachment to ECM biomolecules

We first examined attachment patterns of human endometrial microvascular endothelial cells (HEMECs) on single and pairwise combinations of 10 endometrial-inspired ECM biomolecules (**Fig. 1A**; Collagens I, III, IV, and V, Fibronectin, Laminin, Lumican, Decorin, Hyaluronic Acid, and Decorin) on polyacrylamide gels within the same stiffness regime (∼6 kPa) of the endometrium (∼1-2 kPa)^15–17^. A representative image of the whole microarrays with seeded HEMECs (DAPI stained nuclei) is shown in **Fig. 1B**. Each circular island represents a unique underlying ECM combination. HEMECs display differential attachment patterns as a function of matrix substrate (**Fig. 1C**). Notably, ECM combinations having Collagen I and Collagen IV (blue columns) led to a greater than 3-fold increase in HEMEC attachment (p-value < 0.05) compared to other ECM combinations at 24 hr. Collagen I and Collagen I + Collagen III combinations promoted the highest cellular attachment with a greater than 10-fold increase in attachment compared to Lumican + Tenascin C or Hyaluronic Acid + Tenascin C ECM combinations (**Fig. 1D**).

**Figure 1.**
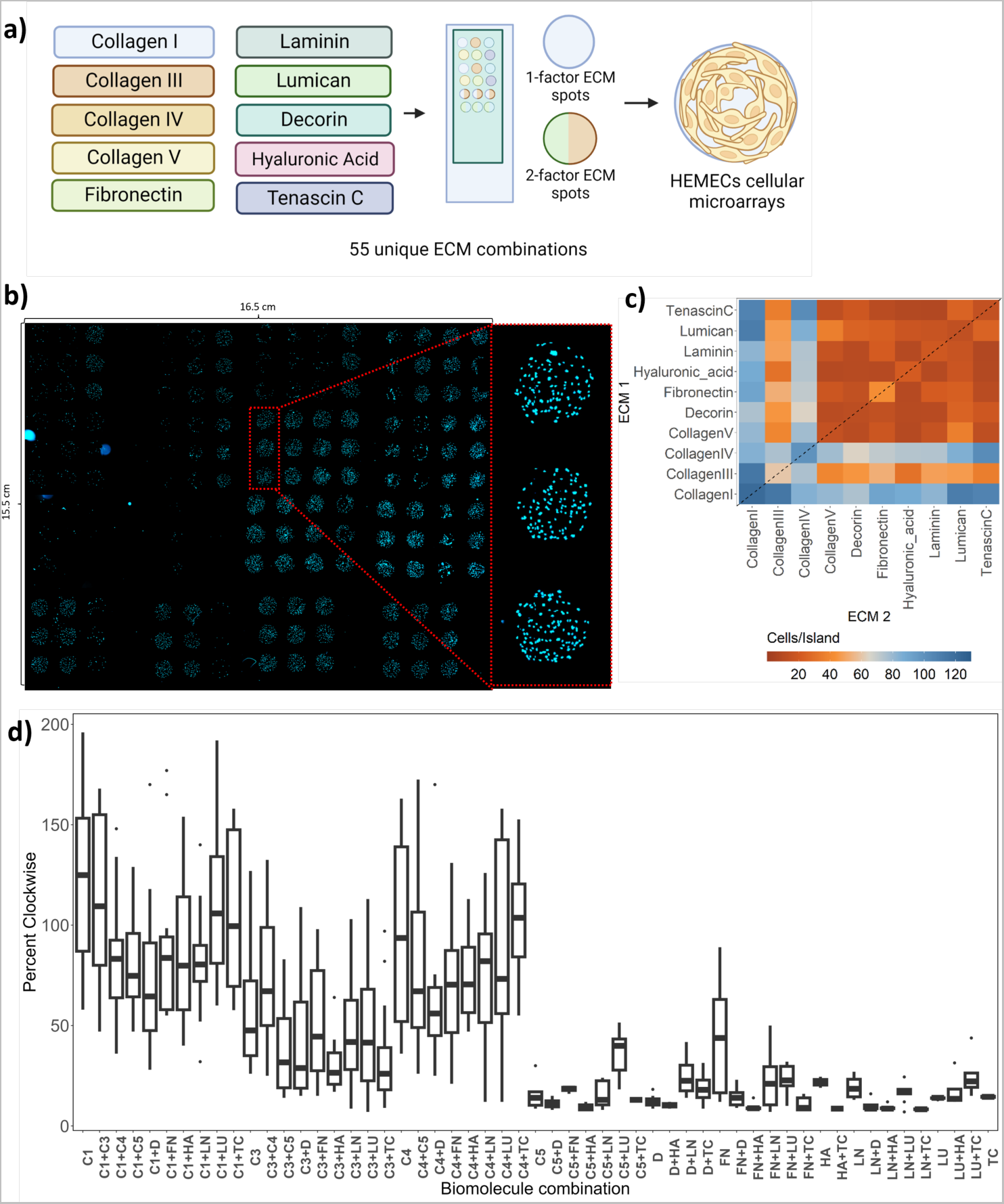
Human endometrial microvascular endothelial cell (HEMEC) attachment on microarrayed extracellular matrix (ECM) combinations. **(A)** Experimental summary. Made in Biorender.com. **(B)** Representative image of HEMECs on islands on microarrays. **(C)** Heat map demonstrating HEMEC/island on ECM combinations. **(D)** Quantification of HEMEC/island on all ECM combinations.

### 2.2. Human endometrial microvascular cells display matrix-induced changes in chirality

We subsequently quantified the influence of matrix biophysical properties on HEMEC chirality via three essential benchmarks of cell chirality on 2D substrates: 1. Stress fiber alignment, 2. Cell boundary-based cell alignment, and 3. Nucleus-centrosome based left-right alignment. In conventional 2D cell culture, the apicobasal cell axis is typically perpendicular to the culture substrate with the front-rear and left-right axes are constrained in plane ^13^. The apicobasal axis is a result of cell attachment to a 2D substrate ^12^. Front-rear chirality is defined along the nuclear-centrosomal axis ^12^. And cytoskeletal chirality can be determined by analyzing the directionality of cellular actin filaments ^18^. Using a high-throughput array-based platform presenting pairwise combinations of same set of 10 endometrial-associated matrix molecules, we observed matrix-induced shifts in the percentage of clockwise actin fibers (**Fig. 2B**). Here, Collagen I (C1) and Collagen I + lumican (C1+LU) promoted the highest percentage of clockwise actin fibers (p-value < 0.05) compared to Collagen III + decorin (C3+D) and Collagen I + laminin (C1+LN) which displayed amongst the lowest clockwise fibers (**Fig. 2C**). Similarly, extracellular matrix ligands altered the percentage of clockwise cells (**Fig. 2E)**. Here, fibronectin and hyaluronic acid (FN+HA) and Collagen III + Laminin (C3+LN) induced the greatest clockwise orientation of HEMEC cells (p-value <0.05) compared to Fibronectin + Decorin (FN+D) and Collagen IV + Fibronectin (C4+FN). We also observed significant matrix-induced shifts in the percentage of left-oriented vs. right-oriented cells based on nucleus-centrosome alignment (**Fig. 2H**). Here, C1 and C1 + LU induced the highest percentage of left-oriented HEMECs compared to HA and Collagen V (C5) which had the lowest percentage of left-oriented cells (**Fig. 2I**). Together, these results demonstrate endometrial endothelial cell chirality is significantly influence by the underlying ECM composition (**Fig. 2**).

**Figure 2.**
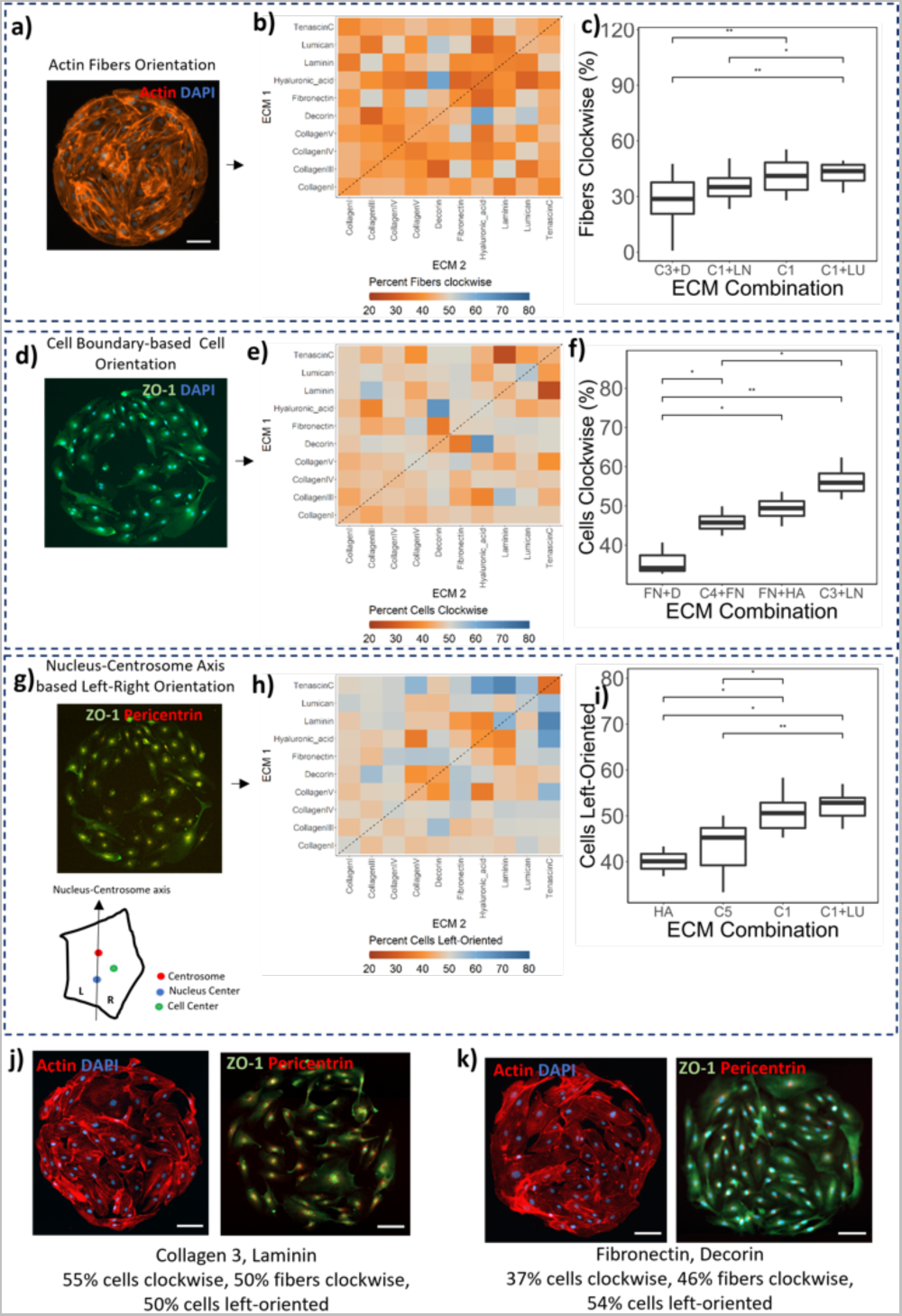
Chirality measurements on microarrays. **(A)** Representative image of actin immunofluorescent staining of human endometrial microvascular endothelial cells (HEMECs) on the microarray extracellular matrix (ECM) island. **(B)** Heat map of average percent clockwise fibers/island on every ECM combination. **(C)** Quantification of actin fiber orientation for specific ECM combinations. **(D)** Representative examples of percent clockwise cells. **(E)** Heat map of average percent clockwise cells/island on every ECM combination. **(F)** Quantification of cell-boundary based orientation on specific ECM combinations. **(G)** Representative Images for calculating nuclear-centrosome axis-based left-right cell orientation. **(H)** Heat map of average percent left-oriented cells/island on specific ECM combinations. **(I)** Quantification of left-right orientation on specific ECM combinations. Representative images of stained HEMECs cultured on **(J)** collagen 3 + laminin and **(K)** fibronectin + decorin. Scale bars: 100 μm.

Interestingly, we observed that shifts in chirality were not always consistent amongst all measured metrics. For example, while the fraction of clockwise cells on C3 + LN substrates (55%) was greater than on FN + D substrates (37%), actin fiber orientation and LR-Orientation of HEMECs were not significantly different between the same matrix combinations (**Fig 2J,K**). Essentially, while HEMECs displayed differential directional rotation properties on C3 + LN and FN + D, they did not exhibit differences in cytoskeletal rotation or alignment across apicobasal and front-rear axis. This suggests that while individual chirality metrics are differentially influenced by the underlying matrix composition, additionally bioinformatics strategies are valuable to understand changes in HEMEC phenotype in a more complex matrix-context dependent manner.

### 2.3. Principal component analysis reveals three distinct chirality-related groups

To understand the multi-dimensional data associated with the chirality of HEMECs as a function of 55 pairwise ECM combinations, we performed principal component analysis (PCA) followed by hierarchical clustering (FactoMineR, R Statistical Computing)^19^. The principal component plane captured 80% variability in the whole data set (**Fig. 3A**), suggesting dimensional reduction significantly retains the original characteristics of the three chirality metrics. To analyze the effect of matrix-induced changes on chirality, we clustered ECM combinations into distinct HEMEC chirality subtypes. We identified three clusters of ECM combinations tied to overall chirality measures (**Fig. 3B-C**). The clusters represented ECM combinations that induced: 1. higher left-oriented cells and lower clockwise cells; 2. higher clockwise fibers and clockwise cells; or 3. lower clockwise fibers and left-oriented cells. Notably, FN + D and LN + TC (tenascin C) substrates were representative of conditions that promoted higher left-oriented cells and lower clockwise cells (Cluster 1). C3 + LN and C5 + FN were representative of conditions that promoted higher clockwise fibers and clockwise cells (Cluster 2). Lastly, C3 + D, FN, and FN + LN were representative of conditions that promoted lower clockwise fibers and left-oriented cells (Cluster 3) (Wilks Test, p-value < 0.05) (**Fig. 3D**). For subsequent experiments, we used these down selected groups of 2-3 representative conditions for each chirality cluster.

**Figure 3.**
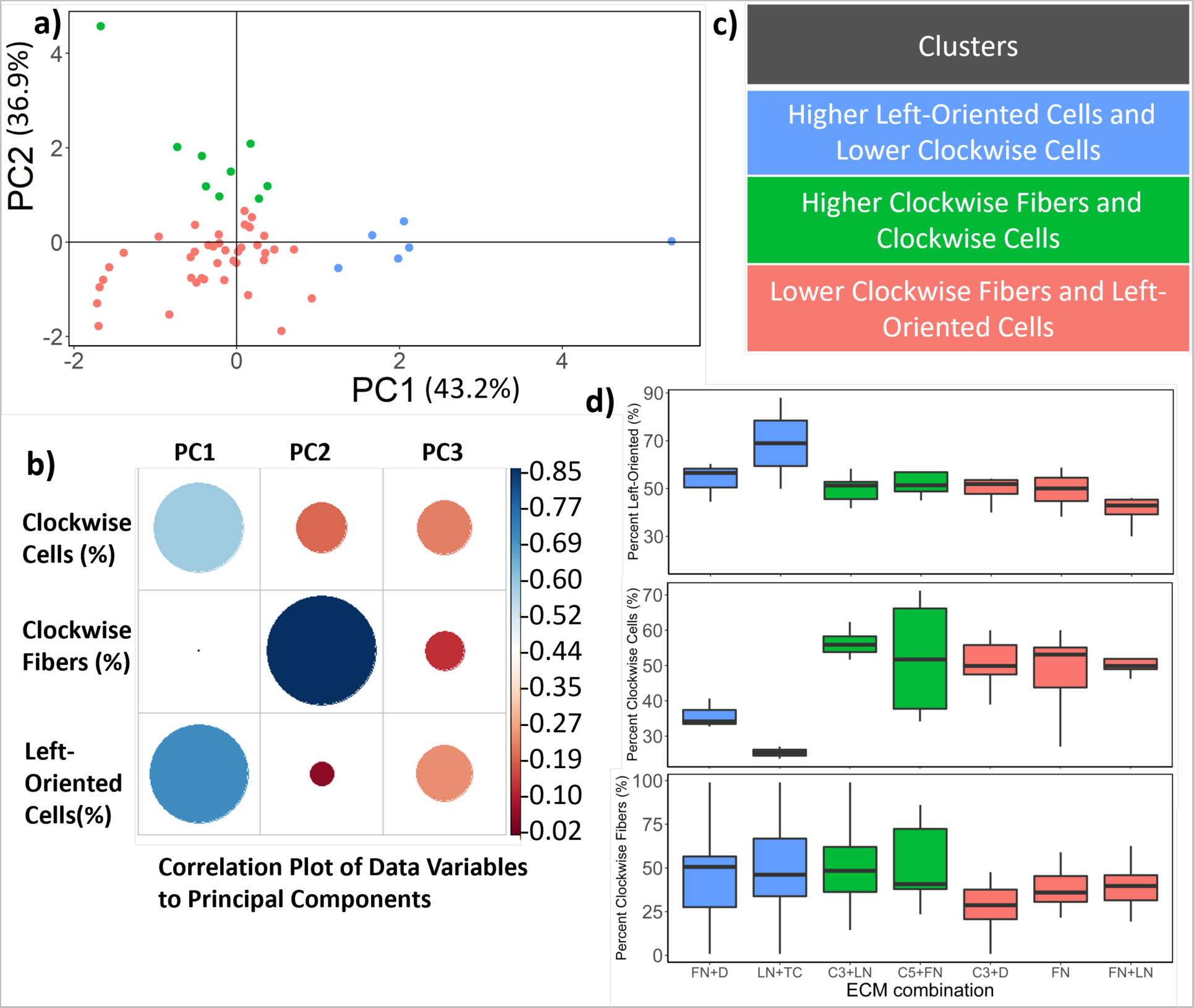
Principal component (PC) analysis of chirality measures. **(A)** Principal component plot of PC1 and PC2. **(B)** Correlation plot of data variables (percentage) to principal components. **(C)** Three distinct clusters revealed from principal component analysis. **(D)** Down selected representative ECM combinations from each cluster.

### 2.4. VEGF and cortisol affect human endometrial endothelial microvascular cell chirality

We subsequently examined whether endometrial-associated biomolecular signals altered patterns of HEMEC chirality cultured on a subset of ECM combinations representing the three chirality clusters identified via PCA (**Fig 3D**, **4A**). The soluble factor conditions were chosen based on endometrial tissue dynamics (**Table 2**) related to angiogenesis and stress: 100 ng/mL vascular endothelial growth factor (VEGF); 10 ng/mL cortisol; 1000 ng/mL cortisol; and control (no exogenous soluble factors)^20^. We quantified chirality fold-change on each ECM combination in response to VEGF/cortisol versus a control (basal) media using the conserved set of chirality metrics: mean cell orientation (clockwise/anti-clockwise/non-chiral); mean actin fiber orientation (clockwise/anti-clockwise/non-chiral); and mean percentage of left/right-oriented cells (**Fig. 4B-D**).

**Figure 4.**
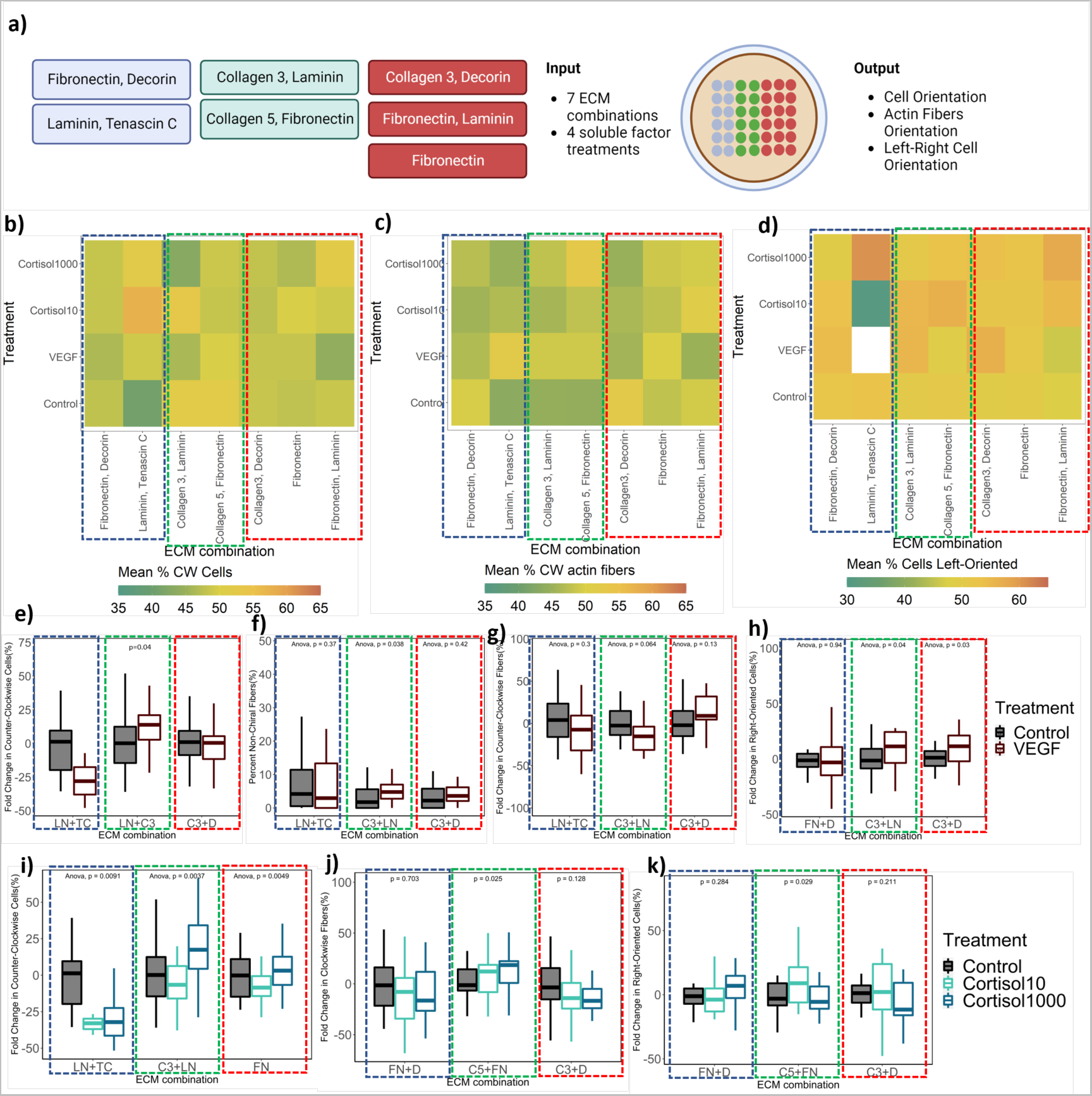
The role of soluble factors on HEMEC chirality. **(A)** Experimental summary. Made inn Biorender.com. Heat maps of soluble factors on **(B)** mean percentage clockwise cells, **(C)** mean percentage clockwise actin fibers, and **(D)** mean percentage left-oriented cells. For Control and +VEGF conditions, **(E)** fold change in counterclockwise cells, **(F)** percent non-chiral fibers, **(G**) fold change in counterclockwise fibers, and **(H)** fold change in right-oriented cells. For Control and + Cortisol Conditions, **(I)** fold change in counterclockwise cells, **(J)** fold change in clockwise fibers, and **(K)** fold change in right-oriented cells. The dashed boxes represent chirality clusters determined in Figure 3.

**Table 1.** Extracellular matrix (ECM) molecules and their relevance to the endometrium.

| **ECM Molecule** | **Location in Endometrium** | **Relevance To Endometrium** | **References** |
| --- | --- | --- | --- |
| Collagen I (C1) | Interstitial fibers; Encapsulate macrostructural elements such as glands and blood vessels | Present during menstrual cycle and pregnancy; Most abundant ECM component in endometrium and decidua | 2,50 |
| Collagen III (C3) | Interstitial fibers; Encapsulate macrostructural elements such as glands and blood vessels | Present during all phases of menstrual cycle; Decreases in decidualization | 2,50 |
| Collagen IV (C4) | Basement membrane and pericellular matrix | Present during menstrual cycle; Increases in decidualization; Deposited by decidual cells to promote trophoblast attachment and invasion | 2,50 |
| Collagen V (C5) | Interstitial fibers; Encapsulate macrostructural elements such as glands and blood vessels | Increases in decidualization; Thought to stabilize growth factors in ECM microenvironment | 2,50 |
| Decorin (D) | Present in decidual ECM | Plays a role in collagen fibrillogenesis but its role in the endometrium is largely unknown | 2,50 |
| Fibronectin (FN) | Basement membrane and pericellular matrix | Abundant in endometrium and decidual ECM; Deposited by decidual cells to promote trophoblast attachment and invasion | 2,50 |
| Hyaluronic Acid (HA) | Stroma and vessels | May influence hydration during the mid-proliferative and mid-secretory phases in the endometrium | 2,50 |
| Laminin (LN) | Basement membrane and pericellular matrix; Decidual tissue | Laminin basement membrane is deposited during reepithelialization, glandular growth, angiogenesis; Deposited by decidual cells to promote trophoblast attachment and invasion; Laminin type varies across cycle and pregnancy | 2,50 |
| Lumican (LU) | Localized to uterine stroma during mice pregnancy; unknown in humans | Unknown in humans | 2,50 |
| Tenascin C (TC) | Localized in stroma; Near stromal cells surrounding proliferating or developing endometrial epithelia; Associated with vascular smooth muscle and is present in late secretory phase spiral arterial walls | May act as inhibitor of cell adhesion to ECM; Found in proliferative phase subepithelial stroma and into early secretory phase in stroma beneath glandular epithelium and absent in secretory phase | 2,50 |

**Table 2.** Hormone and biomolecule conditions.

| **Condition** | **Biomolecules** | **Concentrations** | **Rationale** |
| --- | --- | --- | --- |
| Control | - | Baseline | Baseline control for experimental conditions |
| Vascular Endothelial Growth Factor (VEGF) | VEGF | 100 ng/mL | Positive Control: +VEGF should increase angiogenesis |
| Indolactam | Indolactam | 100 nM | Negative Control: Should decrease angiogenesis |
| Decidualization Condition 1 | Estradiol (E2)<br>Progesterone (P4)<br>Dibutyryl Cyclic AMP (dcAMP) | 10 nM<br>100 nM<br>500 μM | Induces decidualization of endometrial stromal cells |
| Decidualization Condition 2 | 8-Bromo-Cyclic AMP (8-br-cAMP)<br>Medroxyprogesterone Acetate (MPA) | 0.5 mM<br>1 μM | Induces decidualization of endometrial stromal cells |
| Cortisol 1 | Cortisol | 10 ng/mL | Assess stress |
| Cortisol 2 | Cortisol | 100 ng/mL | Assess stress |
| Cortisol 3 | Cortisol | 1000 ng/mL | Assess stress |

Cortisol, but not VEGF, more strongly influenced HEMEC orientation than the underlying ECM conditions. VEGF treatment did not significantly affect chirality on cluster 1 (representing lower clockwise and higher left-oriented cells) ECM combinations. VEGF led to some shifts in chirality for cluster 2 (higher clockwise fibers and clockwise cells), notably a 20%-increase in both the percentage of counter-clockwise cells and right-oriented cells along with a 20%-decrease in the percentage of counter-clockwise actin fibers on C3 + LN (green box) substrates. Soluble VEGF also induced some shifts in chirality for cluster 3 (representing lower clockwise and left-oriented cells), notably a 20% increase in percentage of right-oriented cells (**Fig 4E-H).** Comparatively, cortisol treatment led to a more than 30%-decrease for cluster 1 ECM combination LN + TC as well as a 20% increase in the percentage of counter-clockwise cells for cluster 2 ECM combination LN + FN in a dose-dependent manner (higher clockwise fibers and clockwise cells). Cortisol treatment also led to a significant increase in counter-clockwise cells for cluster 3 ECM FN (**Fig 4I-K**). In contrast to the VEGF treatment, cortisol treatment also lead to a 20% increase in percentage of clockwise fibers on the cluster 2 ECM combination C5 + LN. Importantly, these results suggest that soluble factor cues from the endometrial microenvironment may strongly influence cell chirality, and may have significant implications for considering regarding stress-induced endometrial pathologies (e.g., pre-eclampsia).

### 2.5. Quantification of organizational chirality metrics for HEMEC endothelial tubes

Having established that matrix cues as well as exogenous hormone signals influence the chirality of individual HEMECs, we then examined if these responses extended to the collective behavior of vascular network formation. While endothelial cell chirality has been hypothesized to affect vessel networks^13^, it has not been previously established. Hence, we sought to relate observed shifts in HEMEC cell chirality with endothelial cell vessel network morphology using an *in vitro* Matrigel® tube formation assay which we quantified HEMEC tube network morphology (**Fig. 5A**) in response to 8 different hormone treatments related to endometrial function and angiogenesis (**Table 2**). Hormone conditions included those from HEMEC chirality studies as well as additional factors (medroxyprogesterone acetate, progesterone) related to the decidualization process essential in endometrial vascular maturation and remodeling^20–28^.

**Figure 5.**
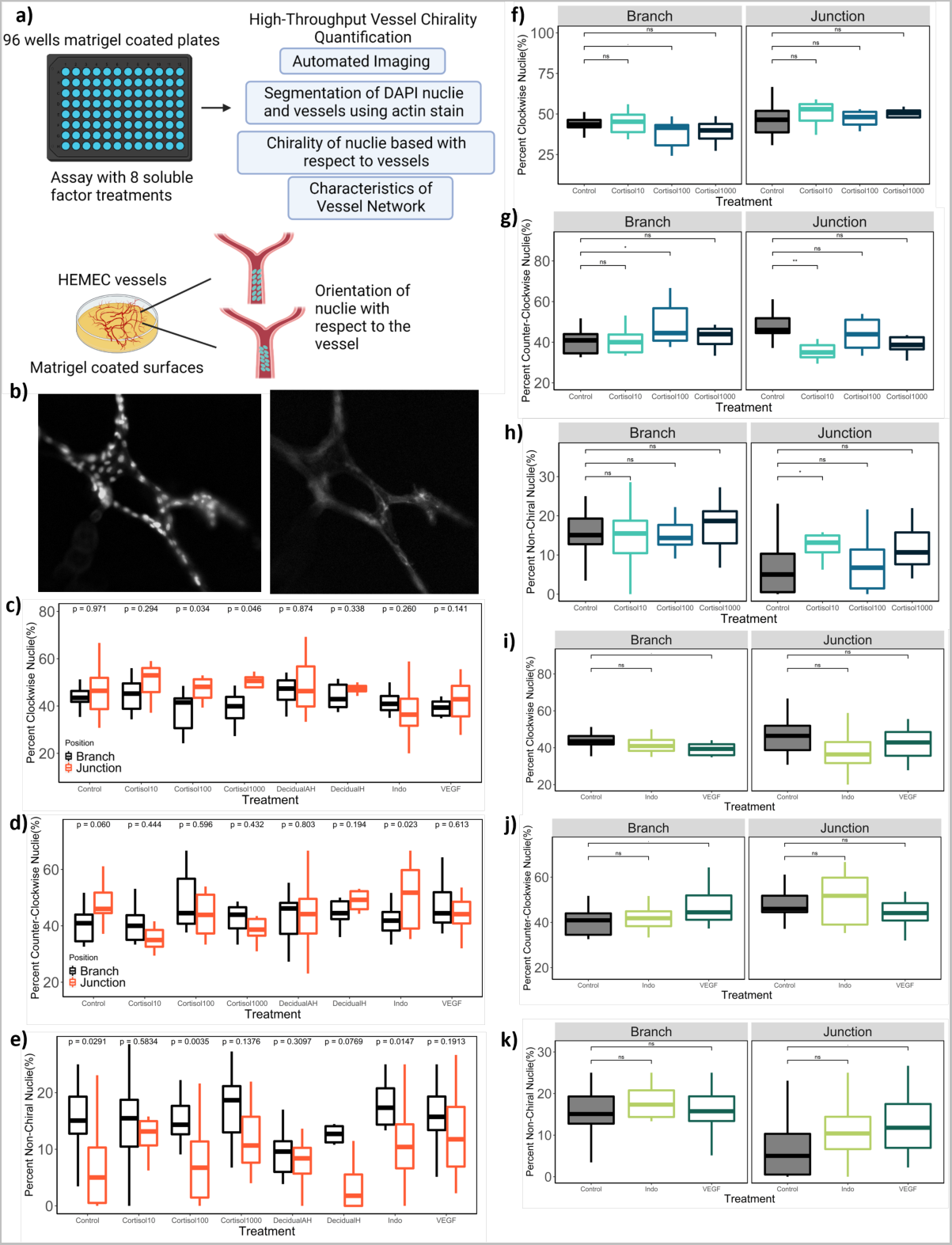
Determination of HEMEC chirality on 96-well phenol red-free Corning Matrigel® Matrix-3D Plate. **(A)** Experimental summary. Made in Biorender.com. **(B)** Representative images of HEMEC tubes stained with DAPI (nuclei) and phalloidin (actin). **(C)** Percent clockwise nuclei in branches and junctions for each soluble factor condition. **(D)** Percent counterclockwise nuclei in branches and junctions for each soluble factor condition. **(E)** Percent non-chiral nuclei in branches and junctions for each soluble factor condition. For branches and junctions in control and +cortisol conditions, **(F)** percent clockwise nuclei, **(G)** percent counterclockwise nuclei, and **(H)** percent non-chiral nuclei. For branches and junctions in control, +indolactam, and +VEGF conditions, **(I)** percent clockwise nuclei, **(J)** percent counterclockwise nuclei, and **(K)** percent non-chiral nuclei.

HEMECs were seeded on Matrigel®-coated 96 well plates and allowed to form endothelial networks for 12 hours under the specific hormone treatments (**Table 2**). The endothelial networks were visualized using actin staining, with cell nuclei visualized using Hoechst (**Fig. 5B**). To quantify the chirality of individual HEMEC cells that formed the endothelial network, the orientation of each Hoechst-stained nucleus was quantified with respect to the endothelial network orientation (**Fig. 5C-E**). Further, regions of each endothelial network were classified into either branch or junction segments based on actin morphology and near-neighbor information. Specifically, a branch endothelial cell was more elongated and had two neighbor cells, whereas junctional endothelial cells had more than 2 neighbors and typically remained more rounded. Interestingly, HEMEC chirality varied for branch vs. junction segments of the endothelial network (**Fig 5F-K**). Junction-associated endometrial endothelial cells displayed significantly higher percentage of clockwise nuclei and in the presence of 100 ng/mL or 1000 ng/mL cortisol (**Fig. 5C**). Interestingly, the percentage of non-chiral nuclei was significantly lower in the junction region except for in the presence of vessel growth and maturation (VEGF and DecidualAH treatment) stimuli, suggesting an overall increase of chiral endothelial cells at network junctions compared to the branches connecting those junctions within endometrial tube network (**Fig 5E**). Inclusion of cortisol drove region-specific shifts in endometrial endothelial cell chirality. Notably, low dose cortisol treatment (10 ng/mL) significantly reduced the percentage of counter-clockwise cells at junctions (increased non-chiral cells vs. control treatment) but the effect at higher concentrations was not significant. However, cortisol treatment at 100 ng/mL reduced the percentage of clockwise cells and increased the percentage of counterclockwise cells (**Fig 5F-H**). These suggest dose-dependent effects of cortisol on HEMEC chirality in different regions of vessel networks. Interestingly, although the anti-angiogenic agent indolactum has been shown to affect chirality of individual endothelial cells^13^, it did not significantly affect nuclei orientation in any region of the HEMEC vessel networks. VEGF treatment did, however, increase the percentage of counterclockwise cells while also decreasing the percentage of clockwise cells, particularly in the stalk (branch) regions connecting junctions (**Fig 5I-K**).

### 2.6. Quantification of HEMEC vessel network characteristics in 2D and 3D

We subsequently examined the effect of HEMEC chirality on metrics of overall endothelial network morphology. We quantified average branch length and branch number of endothelial networks (skeletonized via ImageJ) generated via HEMECs on Matrigel®-coated 96 well plates as a function of hormone exposure (**Fig. 6A**). Exposure to cortisol (100 ng/mL or 1000 ng/mL) significantly decreased average branch length without affecting average number of branches (**Fig 6B-C**) suggesting reduced network complexity. Decreased branch length may be a result of the significant increase in counterclockwise cells in response to 100 ng/mL cortisol treatment (Fig. 5), suggesting changes in cell chirality can induce significant changes in overall vascular network organization and complexity. Further, VEGF treatment (100 ng/mL) induced a decrease in average branch length and slight, but not significant, increase in average branch number compared to untreated control (**Fig 6D-E**) as well as a significant increase in counter-clockwise cells. We subsequently examined if decidualization-associated hormones affected HEMEC networks, chirality, or network architecture. Decidualization is a a hormone-induced process that occurs in the endometrium to form the decidual lining necessary for successful blastocyst implantation by inducing differentiation of endometrial stromal cells. We used two decidualization cocktails, using synthetic progestin (medroxyprogesterone acetate) or progesterone and applied them to the HEMECs as a surrogate for decidualization of stromal cells. Although the average branch length of the resulting HEMEC networks was unchanged between control and decidualized samples (**Fig. 6F**), we noticed HEMEC networks stimulated with progesterone, but not medroxyprogesterone acetate, led to a significant increase in average branch number (and decrease in branch length) as compared to hormone-free controls (**Fig. 6G**). Consistently, we observed significant changes in cell chirality in the presence of the stress hormone cortisol for both microarrays and HEMEC network architecture assays.

**Figure 6.**
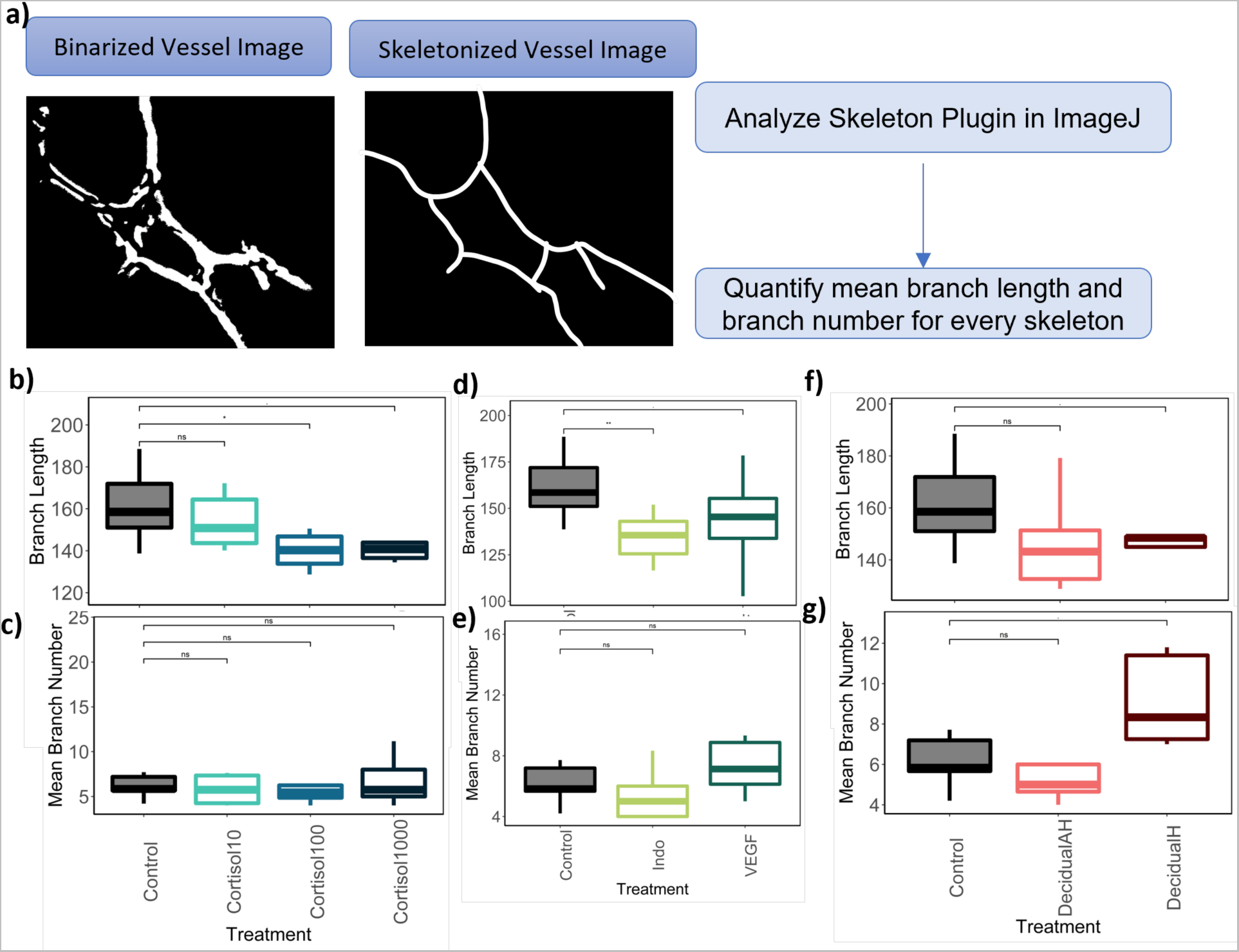
Quantification of mean branch length and branch number across conditions on 96-well phenol red-free Corning Matrigel® Matrix-3D Plate. (A) Experimental summary. Made in Biorender.com. For control and +cortisol conditions, **(B)** mean branch length (pixel number) and **(C)** mean branch number. For control, +indolactam, and +VEGF conditions, **(D)** mean branch length (pixel number) and **(E)** mean branch number. For control and +hormone conditions, **(F)** mean branch length (pixel number) and **(G)** mean branch number.

We subsequently co-cultured HEMECs with human endometrial stromal cells in a methacrylamide-functionalized gelatin (GelMA) hydrogel in a manner previously reported to generate endometrial perivascular niche cultures^20^ to more closely assess cortisol-associated changes in three-dimensional endothelial network architecture (**Fig. 7A**). We did not observe significant changes in total network length/mm^3^, total branches, or total vessels in response to increasing cortisol concentration(**Fig. 7B-E**); however, the 1000 ng/mL concentration decreased branch length compared to all other conditions (**Fig. 7D**). Importantly, these results are consistent with the decrease in branch length observed using Matrigel® in the presence of 1000 ng/mL cortisol **(Fig. 6B-C),** the significant increase in clockwise cells at HEMEC network junctions **(Fig. 5C**), and matrix-dependent HEMEC chirality changes in response to 1000 ng/mL cortisol (**Fig. 4I-K**). Together, these results demonstrate that cortisol increases HEMEC cell chirality in a matrix dependent manner, more strongly affects HEMEC chirality at discrete points (junctions) within 3D endothelial networks, and induces the formation of denser, more compact three-dimensional endometrial networks in gelatin hydrogels.

**Figure 7.**
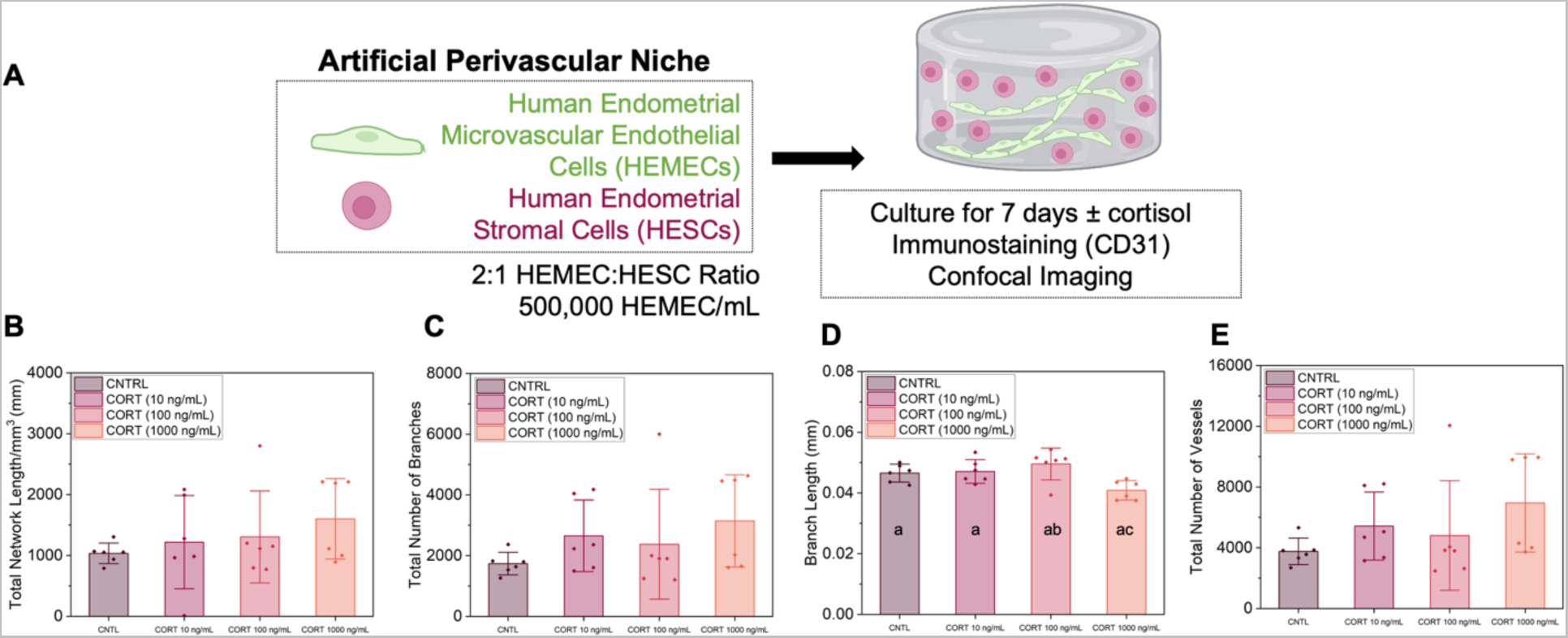
The effects of cortisol on endometrial perivascular niche complexity. **(A)** Experimental summary. **(B)** Quantification of total vessel length per mm^3^, **(C)** total number of branches, **(D)** average branch length, and **(E)** total number of vessels for control and cortisol treated samples (n=6 hydrogels per condition; 3 ROI imaged per gel and averaged). Groups with different letters are statistically significantly different from each other. Data presented as mean ± standard deviation. Created with Biorender.com.

## 3. Discussion

In this work, we defined the influence of extracellular matrix ligands and soluble molecules associated with the endometrial tissue microenvironment on the chirality of human endometrial microvascular endothelial cells (HEMECs). Collagens I-VI have a profound effect on HEMEC attachment on 2D microarray studies. Specifically, collagens I-IV alone or in combination with other ECM biomolecules increased HEMEC cell attachment on 2D microarrays compared to matrix combinations with Collagen V and the other endometrial associated ECM biomolecules (fibronectin, laminin, lumican, decorin, hyaluronic acid, tenascin C). Interestingly, two combinations we previously showed to increase endometrial epithelial cell (EEC) attachment (Collagen I + Collagen III, Collagen IV + Tenascin C)^14^ also promoted high levels of HEMEC attachment, suggesting these combinations may be more universally important in the development of artificial endometrial models. Notably, collagen I alone resulted in the highest average HEMEC attachment per island which is not surprising because collagen I is the most prevalent collagen in the endometrium^2^.

We then sought to define the role of the matrix biomolecular environment on HEMEC chirality via three measures of cell chirality on microarrays: 1. Actin fiber orientation, 2. Cell boundary-based cell orientation, and 3. Nucleus-centrosome axis-based left-right orientation.

Micropatterned surfaces have been shown to be accurate and reproducible systems for quantifying cell chirality in vitro^12, 29^. Although prior work showed that chirality is relevant to most cells and is phenotype-specific,^12^ prior work considering endothelial cells have largely used human umbilical vein endothelial cells (HUVECs) cultured on fibronectin-coated wells to investigate links between chirality and endothelial cell permeability^13^. Those studies showed predominantly rightward bias of endothelial cells (HUVECs, mouse vena cava, mouse thoracic aorta, mouse aortic arch). Here, we report a predominantly leftward and counterclockwise bias for endometrial endothelial cell on fibronectin. Future work warrants exploring whether there are additional functional differences in endometrial endothelial cells, such as permeability and tight junction formation, based on chirality.

Unlike previous studies which largely defined endothelial cell chirality on only fibronectin, we characterized endometrial endothelial chirality across 55 ECM combinations of 10 different ECM biomolecules. Our work significantly advances methodology used to characterize chirality and shows endometrial endothelial cell chirality can be influenced the extracellular matrix signals. To our knowledge, this work is the first to systematically study chirality across multiple ECM biomolecules in a high-throughput manner. We found that HEMEC actin fibers and cells largely displayed counterclockwise biases across the different ECM conditions, but the left-right cell orientation was more variable in response to matrix ligand presentation. These results suggest that although endometrial endothelial cells largely prefer a counterclockwise orientation at the cell-scale, their internal orientation is sensitive to the underlying matrix condition on which they are cultured. We then identified three primary clusters of endometrial endothelial cell chirality metrics via principal component analysis, Namely, HEMECs that were: 1. Higher left-oriented and lower clockwise-oriented, 2. Higher clockwise oriented with increased clockwise actin fibers, and 3. Reduced left-oriented and with fewer clockwise actin fibers. We down selected 2-3 conditions per principal component cluster to study further.

We then showed soluble biomolecules from the endometrial tissue microenvironment were sufficient to alter endothelial cell chirality established by matrix signals. For these studies, we examined three factors commonly associated with the endometrial microenvironment: VEGF and 10 or 1000 ng/mL of the stress hormone cortisol. These conditions were chosen based on concentrations we had previously shown to influence endometrial-trophoblast interactions in engineered endometrial tissue mimics^20, 30, 31^ Overall, these data suggested that endometrial endothelial cell chirality is jointly influenced by matrix and biomolecule signals, with cortisol most strongly affecting counterclockwise cell orientation across all three chirality cluster conditions.

We subsequently examined chirality within multicellular endometrial endothelial networks. Endothelial cells do not exist as single cell entities, but rather as multicellular entities that define endometrial vessel networks. This is particularly acute in the endometrium, where significant vascular regression and regrowth occurs rapidly across the menstrual cycle. We adapted a automated analysis pipeline^20, 32^ to quantify chirality within endothelial networks using a cell-based chirality metric. We used segmentation and object detection to identify endothelial branches versus junctions within HEMEC networks, then defined nuclei orientation with respect to the local vessel network orientation. Using this method, we characterized shifts in endometrial endothelial cell chirality in response to 5 discrete soluble factors associated with the endometrial tissue microenvironment. Excitingly, shifts in individual cell chirality were not only dependent on soluble factors but also on the relative (along a branch vs. at a junction) position of the endometrial endothelial cell within a larger endothelial network. For example, 10 ng/mL cortisol affected the chirality of the HEMECs only at junctions whereas 100 ng/mL cortisol affected HEMEC chirality along branches and reduced the average branch length of the endothelial networks. Junctions are a particularly significant concept in the formation and remodeling of branched endometrial endothelial networks over the course of rapid vascular remodeling that occurs across the menstrual cycle.

Although we performed the high-throughput endothelial cell network architecture studies using a commonly used Matrigel® tube formation assay, a significant limitation of Matrigel® is heterogeneous in terms of its composition^33, 34^, so it is possible its batch-to-batch variability may affect cell chirality measures. To address this limitation, we adapted a three-dimensional endothelial cell network assay to quantify endometrial networks in a material developed from only a single ECM molecule(methacrylamide-functionalized gelatin, GelMA). We examined the explicit role of cortisol on three-dimensional endothelial cell networks in GelMA hydrogels because cortisol most strongly impacted endometrial endothelial cell chirality in both 2D microarrays and in Matrigel®. We found that addition of 1000 ng/mL of cortisol impacted the average branch length of the endothelial networks, suggesting that cortisol plays an important role in regulating macro cellular structures formed by endometrial endothelial cells. Taken together, this work demonstrates that exogenous cortisol strongly influences endometrial cell chirality at the individual cell level, influences cell chirality as a function of position within endometrial endothelial cell networks, and strongly regulates the overall complexity of endometrial endothelial cell networks (e.g., vessel branch length), with results conserved between multiple experimental platforms in both two-dimensions and three-dimensions.

In summary, we report a comprehensive approach that defined the chirality of human endometrial endothelial cells in response to endometrial-associated ECM matrix and soluble biomolecule conditions. Notably, endometrial endothelial cells are chiral, with their chirality strongly dependent on both underlying matrix and inflammatory biomolecules conditions. Further, their chirality is highly contextual within larger cellular cohorts, with unique responses observed along endothelial branches versus at junctions within endothelial networks. Our findings have implications in endometrial physiology and for a range of endometrial disorders. Notably, changes in endometrial ECM composition observed during disease progression or in response to abherrent remodeling could alter endometrial cell chirality and ultimately effect vessel network remodeling and stability. For endothelial cells, this would be especially important for vascular-related diseases and disorders such as endometriosis or preeclampsia, where insufficient vascular remodeling in the first trimester detrimentally impacts placental function through the entirety pregnancy^35, 36^. Importantly, we also demonstrate the stress-associated factor cortisol significantly impacts endometrial endothelial network formation and cell chirality. This work could inform creation of therapeutics and molecular targets to mitigate disorders related to the endometrial vascular and suggests that stress may play important roles in the menstrual cycle and endometrial function.

## Online Methods

### 2.1. Cell Culture

#### 2.1.1. Human Endometrial Microvascular Endothelial Cell Culture

Human endometrial microvascular endothelial cells (HEMEC; ScienCell Catalog #7010) were maintained as per the manufacturer’s instructions. Cell were cultured in phenol red-free Endothelial Cell Medium (ECM; ScienCell Catalog #1001-prf) supplemented with an endothelial cell growth supplement (ScienCell Catalog #1052), 5% charcoal-stripped fetal bovine serum (Sigma-Aldrich F6765), and 1% penicillin/streptomycin (Invitrogen 15140-122). We used charcoal-stripped fetal bovine serum to reduce the steroid hormone concentrations in the cell medium as to not cofound the data. HEMECs were used experimentally no more than 5 passages from purchase and were cultured in 5% CO_2_ incubators at 37°C. Routine mycoplasma testing was performed on the cells using a MycoAlert^TM^ Mycoplasma Detection Kit (Lonza). Cell ancestry information (e.g., racial and ethnic background, age, gender identity) was not provided by the vendor although the cell ancestry may affect cellular behavior and response ^37, 38^.

#### 2.1.2. Human Endometrial Stromal Cells

Human endometrial stromal cells (HESC; ATCC® CRL-4003) were maintained as per the manufacturer’s instructions. Cells were cultured in custom phenol red-free DMEM/F-12 (based on Sigma #D 2906) from the SCS Cell Media Facility supplemented with 1% ITS+ Premix (Corning 354352), 500 ng/mL puromycin (Millipore Sigma P8833), 10% charcoal stripped fetal bovine serum (Sigma-Aldrich F6765), and 1% penicillin/streptomycin. HESC were used experimentally no more than 5 passages from purchase and were cultured in 5% CO_2_ incubators at 37°C. Routine mycoplasma testing was performed using a MycoAlert^TM^ Mycoplasma Detection Kit (Lonza). Cell ancestry information (e.g., racial and ethnic background, age, gender identity) was not provided by the vendor although the cell ancestry may affect cellular behavior and response ^37, 38^.

#### 2.1.3. Hormone and Biomolecule Culture Treatments

Table 2 describes each hormone and biomolecule treatment cocktail in this work. The hormones and biomolecules were diluted in media containing charcoal-stripped fetal bovine serum. The biomolecules chosen were pro-angiogenic (± 100 ng/mL recombinant human VEGF_165_; PeproTech 100-20), anti-angiogenic (Indolactam; Sigma Aldrich I0661), representative of stress (cortisol 10-1000 ng/mL; Sigma Aldrich H0888), or representative of stromal cell decidual response. Decidualization was induced by adding one of the following hormone cocktails to cell media: (i) synthetic progesterone based, 1 μM medroxyprogesterone acetate (MPA; Sigma-Aldrich M1629) + 0.5 mM 8-bromodenosine 3’,5’-cyclic monophosphate (8-Br-cAMP; Sigma-Aldrich B5386) or (ii) progesterone based, 0.5 mM dibutyryl cyclic AMP (dcAMP; Millipore Sigma 28745) + 10 nM estradiol (E2; Sigma-Aldrich E2758) + 100 nM progesterone (P4; Sigma-Aldrich P8783).

### 2.2. Microarray Fabrication and Assays

#### 2.2.1. Microarray Fabrication

Polyacrylamide (PA) hydrogels were prepared following previous protocols^16, 17, 39^. Briefly, 12 mm glass coverslips and standard microscopy glass slides were etched by immersing them 0.2 N NaOH (Sigma-Aldrich 415413-1L) for 1 hour on an orbital shaker and then rinsing with dH2O. The coverslips were then air-dried and placed on a hot plate at 110°C until dry. For silanization, the cleaned coverslips and slides were immersed in 2% v/v 3-(trimethoxysilyl)propyl methacrylate (Sigma Aldrich 440159-500ML) in ethanol and placed on the shaker for 30 minutes, followed by a wash in ethanol for 5 minutes. The silanized coverslips and glass slides were air-dried, and again placed on the hot plate at 110°C until dry. For fabrication of hydrogels with specific elastic moduli, prepolymer solution in dH20 with 8% acrylamide (Sigma-Aldrich A3553-100G) and 0.55% bis-acrylamide (Sigma-Aldrich M7279-25G) was prepared to achieve elastic moduli of 6 kPa. The prepolymer solution was then mixed with Irgacure 2959 (BASF, Corp.) solution (20% w/v in methanol) at a final volumetric ratio of 9:1 (prepolymer:Irgacure).

This working solution was then deposited onto Rain-X (Amazon Rain-X 800002245) coated slides (20uL/coverslip) and covered with silanized coverslips. The sandwiched working solution was transferred to a UV oven and exposed to 365 nm UV A for 10 min (240E3 µJ). The coverslips with the hydrogels attached to it were immersed in dH2O at room temperature for a day in order to remove excess reagents from the hydrogel substrates. Before microarray fabrication, hydrogel substrates were thoroughly dehydrated on a hot plate for ≥15 minutes at 50°C. Microarrays were fabricated as described previously [37-39]. ECMs (Table 1) for arraying were diluted in 2×ECM printing buffer. to a final concentration of 250 μg/mL and loaded in a 384-well V-bottom microplate. To prepare 2× ECM protein printing buffer, 164 mg of sodium acetate and 37.2 mg of ethylenediaminetetraacetic acid (EDTA) was added to 6 mL dH2O. After solubilization, 50 µL of pre-warmed Triton X-100 and 4 mL of glycerol was added. 40 – 80 µL of glacial acetic acid was added, titrating to adjust the pH to 4.8.A robotic benchtop microarrayer (OmniGrid Micro, Digilab) loaded with SMPC Stealth microarray pins (ArrayIt) was used to microprint ECM combinations from the 384 microwell plate to polyacrylamide hydrogel substrate, resulting in ∼600 µm diameter arrayed domains. Fabricated arrays were stored at room temperature and 65% RH overnight and left to dry under ambient conditions in the dark.

#### 2.2.2. Microarray Cell Seeding

Microarrays were seeded as previously described ^40–43^. Microarrays were sterilized in 1% penicillin/streptomycin in PBS under UV light for 20 minutes. HEMECs were trypsinized and seeded onto each microarray by dividing the total amount of trypsinized cells equally amongst the arrays (approximately 200,000-500,000 cells/slide arrays-55 ECMs and 10,000-50,000 per coverslip array -7 ECMs). The seeding density was maintained to get confluent islands and kept constant for different conditions in each biological replicate. The arrays were gently shaken every 30 minutes for 2 hours and subsequently rinsed once after 6 hours with cell medium to remove any unattached cells. Cells were subsequently fixed 24 hours after the wash step.

### 2.3. Matrigel® Tube Formation Assays

As per the manufacturer’s instructions, a 96-well phenol red-free Corning Matrigel® Matrix-3D Plate (Corning 356259) was prepared for culture and seeded with 10,000 HEMEC/well. Sample well replicates (n=6-8) were prepared for each condition. Samples were fixed 12 hours after seeding and subsequently stained, imaged, and analyzed.

### 2.4. Endometrial Perivascular Niche

Endometrial perivascular niche cultures were fabricated as described previously by co-culturing HEMEC and HESC in methacrylamide-functionalized gelatin (GelMA) hydrogels^20^. We utilized a previously characterized batch of GelMA with a degree of functionalization of 57%, determined via ^1^H-NMR ^30, 44, 45^. First, we sterilized lyophilized GelMA for 30 minutes under UV light. We then made hydrogel pre-polymer solution using a solution consisting of lyophilized GelMA (5 wt%) dissolved at 37°C in phosphate buffered saline (PBS; Lonza 17-516F) and combined with 0.1% w/v lithium acylphosphinate (LAP) as a photoinitiator. HEMEC and HESC were then passaged and combined in a 2:1 endothelial to stromal cell ratio with a concentration of 500,000 HEMEC/mL. Cell-laden pre-polymer solution was polymerized under UV light (λ=365 nm, 7.14 mW cm^-2^; AccuCure Spot System ULM-3-365) for 30 s to create hydrogels. Cell laden hydrogels were maintained in 48 well plates for 7 days and maintained in 800 μL ECM with or without cortisol. Medium was replaced every 3 days. This experiment used charcoal-stripped fetal bovine serum (Sigma-Aldrich F6765) instead of regular fetal bovine serum to decrease endogenous hormones in the base ECM.

To quantify vessel network metrics, hydrogels were fixed on day 7 and subsequently stained with CD31 using previously published protocols^20^. Briefly, hydrogel samples were permeabilized for 15 minutes in a 0.5% Tween20 (Fisher Scientific BP337) solution, washed 3x5 minutes in 0.1% Tween20 solution (PBST), blocked for 1 hour at room temperature in a 2% Abdil solution (2% bovine serum albumin; Sigma Aldrich A4503 + 0.1% Tween20), and subsequently incubated in primary antibody solution (1:200 CD31 Dako IS610) overnight at 4°C. The following day, 4x20 minutes washes with PBST were performed and then hydrogel samples were incubated with secondary antibody solution (1:500 Alexafluor 488 goat anti-mouse Thermo Fisher A-11001) overnight at 4°C. Hydrogel samples were then washed 4x20 minutes with PBST, incubated for 30 minutes in Hoechst (1:2000; Thermo Fisher H3570), washed a final time in PBST, and were stored in PBST until imaged. Hydrogel samples were imaged using glass bottom confocal dishes (In Vitro Scientific, D29-20-1-N) on a DMi8 Yokogawa W1 spinning disc confocal microscope outfitted with a Hamamatsu EM-CCD digital camera (Leica Microsystems). We collected three 100 μm z-stacks with a 5 μm step size for each hydrogel for 3 regions of interest (ROI). We then used a computational pipeline consisting of a FIJI macro and custom MATLAB algorithm ^32, 46^ to quantify total network length / mm^3^, average branch length (network length / number of vessels), number of branches, and number of vessels for each sample as previously described^20^.

### 2.5. Chirality Immunostaining

Samples were stained for ZO-1 (5 μg/mL; Thermo Fisher 40-2200) and pericentrin (1 μg/mL; Abcam ab28144) or phalloidin (Actin; Abcam ab176758). Fixed microarray or Matrigel samples were permeabilized with 0.5% Tween20 in PBS for 15 minutes at room temperature followed by 3x5 minute washes in PBST (0.1% Tween20 in PBS). Samples were subsequently blocked in 2% abdil solution (bovine serum albumin, Tween20, and PBS) for 1 hour at room temperature. For ZO-1 and pericentrin stained samples, primary antibody solution was added to samples followed by overnight incubation at 4°C. After primary antibody incubation, samples were washed 3x5 minutes with PBST and subsequently incubated in secondary antibody solution (1:500; Alexafluor 488 goat anti-rabbit Thermo Fisher A-11008 or Alexafluor 555 goat anti-mouse Thermo Fisher A-21422) overnight at 4°C. Following secondary antibody incubation, samples were washed 3x5 minutes in PBST and mounted with DAPI Fluoromount (microarrays) or stained for 30 minutes using Hoechst followed by a PBST wash (Matrigel).

For actin-stained samples, all steps outlined above were performed. After the block step, an anti-actin antibody was diluted 7 μL antibody to 1000 μL blocking solution and samples were incubated in this solution for 1 hour at room temperature followed by 3 quick PBS washes.

### 2.6. Microscopy

The microarrays for the immunostaining were imaged using Axioscan.Z1 Slide Scanner and 10X objective. A wide tile region was defined for the whole array region which was then stitched offline using Zen and exported into TIFF Images for each individual channel. Images of entire arrays were converted to individual 8-bit TIFF files per channel (i.e., red, green, blue) by Fiji (ImageJ version 1.52p) [41]. The images were cropped in MATLAB (version R2018b) to separate each array in a single image. Positional information for each array was automatically calculated using their relative position from the positional dextran-rhodamine markers. CellProfiler (version 4.0.0) [42] was used get per cell measurement for each channel. Nuclei were identified using the DAPI channel image using IdentifyPrimaryObject module and other stains were associated with a specific nucleus was identified by looking at the red/green stain around these nuclei using IdentifySecondaryObject module. The MeasureObjectIntensity module was used to quantify single-cell intensity. The data were exported to CSV files that were then imported in RStudio for data visualization.

### 2.7. Chirality Analysis

We quantified measurements using automated cell segmentation (Identify Primary and Secondary Objects) via Cell Profiler^47^ followed by radial alignment calculation of each cell on an ECM island using Rstudio. For all chirality quantification, cell-based angle and coordinated were automatically obtained using custom pipelines made in Cell Profiler and analyzed further in RStudio. For cellular orientation, ZO-1 stain was used to detect cellular boundary and object detection. Using MeasureObjectSizeandShape module in Cell Profiler, Orientation of each cell with respect to the x-axis of the image was obtained. This orientation was converted to radial orientation with respect to the centroid of the circular island using a RScript. For Actin Fiber Orientation, the fibers were segmented and detected using object detection using Cell Profiler. As before, radial orientation of the fibers were calculated RScript. For Left-Right Orientation, DAPI object (nucleus) center (Primary Object), ZO-1 object (Cellular) center (Secondary Object) and Pericentrin Center (Tertiary Object related to Secondary Object) was detected using Cell Profiler. The Left-Right Orientation was calculated based on these three coordinates for each cells on an island using an RScript.

For chirality quantifications on the 3D vessel networks, actin stain was used to segment the vessel objects and orientation was obtained. Independently DAPI objects were segmented and orientation was obtained. The DAPI objects and actin objects were related to each other based on spatial overlap. This meant that each DAPI nucleus was related to its subsequent vessel object, which important for quantification of relative orientation of DAPI nucleus with respect to the vessel. The branch and junction characterization of vessel region was done using a filter on vessel ellipticity and number of neighbors.

### 2.8. Statistics

All microarray experiments consisted of at least three biological replicates, with at least 15 technical replicates (islands or microwells) per biological replicate per combination of treatment and readout. For comparison between conditions in this study, Wilcoxon tests were performed using the Wilcox.test function in R. *P* values of < 0.05 were considered significant. HEMEC-HESC PVN data were analyzed using OriginLab 2021b and RStudio for n=6 hydrogels per condition. Normality and homoscedasticity was determined via Shapiro-Wilkes and Levene’s test, respectively. Based on these results, we ran Kruskall-Wallis ANOVA and Dunn’s post hoc test (non-normal, homoscedastic). Data are presented as mean ± standard deviation with significance set to *p* < 0.05.

## Acknowledgements

Research reported was supported by the National Institutes of Diabetes and Digestive and Kidney Diseases of the National Institutes of Health under Award Numbers R01 DK0099528 (B.A.C.H.) and R01 DK125471 (G.H.U.), the National Cancer Institute of the National Institutes of Health under Award Number R01 CA256481 (B.A.C.H), and by the National Institute of Biomedical Imaging and Bioengineering of the National Institutes of Health under Award Number T32 EB019944 (S.G.Z.). The content herein is solely the responsibility of the authors and does not necessarily represent the official views of the National Institutes of Health. The authors also gratefully acknowledge additional funding provided by the Department of Chemical & Biomolecular Engineering and the Carl R. Woese Institute for Genomic Biology at the University of Illinois Urbana-Champaign. The authors also thank the Institute for Genomic Biology Core Facilities (Dr. Austin Cyphersmith) at the University of Illinois Urbana-Champaign for assistance with imaging and Dr. Cody Crosby and Dr. Janet Zoldan for providing the image analysis pipeline for endothelial network characterization.

## Author Contributions

We describe contributions to the manuscript using the Contributor Roles Taxonomy (CRediT) ^48, 49^: *Writing – Original Draft*: SGZ and IJ; *Writing – Review & Editing:* SGZ, IJ, GHU, and BACH; *Conceptualization:* SGZ and IJ; *Investigation:* SGZ, IJ, and HT; *Methodology:* SGZ and IJ; *Formal Analysis:* IJ; *Data Curation:* SGZ and IJ; *Visualization:* SGZ and IJ; *Project Administration:* BACH and GHU; *Resources:* GHU, and BACH; *Funding Acquisition*: GHU and BACH; Supervision: GHU, and BACH.

## Data Availability

The raw data required to reproduce these findings are available per request by contacting the corresponding author. The processed data required to reproduce these findings are available per request by contacting the corresponding author.

